# Macaque anterior cingulate cortex deactivation impairs performance and alters lateral prefrontal oscillatory activities in a rule-switching task

**DOI:** 10.1101/427302

**Authors:** Liya Ma, Jason L. Chan, Kevin Johnston, Stephen G. Lomber, Stefan Everling

## Abstract

In primates, both the dorsal anterior cingulate cortex (dACC) and the dorsolateral prefrontal cortex (dlPFC) are key regions of the frontoparietal cognitive control network. To study the role of the dACC and its communication with the dlPFC in cognitive control, we recorded the local field potentials from the dlPFC before and during the reversible deactivation of the dACC, in macaque monkeys engaging in uncued switches between two stimulus-response rules. Cryogenic dACC deactivation impaired response accuracy during rule-maintenance, but not rule-switching, which coincided with a reduction in the correct-error difference in dlPFC beta activities specifically during maintenance of the more challenging rule. During both rule switching and maintenance, dACC deactivation prolonged the animals’ reaction time and reduced task-related theta/alpha activities in the dlPFC; it also weakened dlPFC theta-gamma phase-amplitude modulation. Thus, the dACC and its interaction with the dlPFC plays a critical role in the maintenance of a new, challenging rule.

## INTRODUCTION

Survival in a dynamic environment requires cognitive control, which is the brain’s ability to guide actions using relevant information in a given context while suppressing irrelevant input, and to flexibly adjust such guidance when the context changes. As part of the primate cognitive control network, the dorsolateral prefrontal cortex (dlPFC) and the dorsal anterior cingulate cortex (dACC) have been shown to co-activate in various cognitively demanding tasks (Brown et al., 2007; Cabeza and Nyberg, 2000; Dosenbach et al., 2008; Dosenbach et al., 2006; Duncan and Owen, 2000). Although the literature suggests that the dACC monitors and evaluates the cost and benefits of controlled actions (Shenhav et al., 2013) and environments (Kolling et al., 2012), while the dlPFC allocates and regulates the control needed to execute the chosen action (Botvinick et al., 2001; Holroyd and Yeung, 2012), their functions often overlap. Similar to the dACC, the dlPFC also encodes reward expectation (Kobayashi et al., 2006; Wallis and Miller, 2003; Watanabe, 1996) and prediction error (Glascher et al., 2010). Hence it is likely that the two regions interact closely during cognitive control processes (Alexander and Brown, 2015), sharing information concerning contexts, actions, outcomes and their values. Thus, understanding the mechanisms through which the two areas communicate will shed light on the neural processes of cognitive control.

To study the role of the dACC in cognitive control processes such as behavioral flexibility and the inhibition of irrelevant task rule, we trained macaque monkeys on uncued switches between two stimulus-response rules—looking either toward (prosaccade) or away from (antisaccade) a peripheral visual stimulus. We hypothesize that dACC deactivation would impair the animals’ performance on the task, as it was found to encode rule information well before the onset of the peripheral target stimulus (Johnston et al., 2007; Womelsdorf et al., 2010). Additionally, given dACC’s involvement in feedback processing (Amiez et al., 2005; Holroyd and Coles, 2002; Kerns et al., 2004; MacDonald et al., 2000) and prediction error (Alexander and Brown, 2010; Hyman et al., 2017), dACC deactivation may result in perseveration and a delay to the rule switch. Alternatively, dACC was suggested to be critical for sustaining effective choices based on reward history (Chudasama et al., 2013; Kennerley et al., 2006), which predicted an impairment in maintaining performance on the new rule but not a delay in rule switching *per se*. Lastly, the dACC’s role in controlling cognitive effort (Chong et al., 2017; Croxson et al., 2009; Klein-Flugge et al., 2016; Parvizi et al., 2013; Rudebeck et al., 2006; Vassena et al., 2014; Walton et al., 2003) predicts a deficit in post-switch performance maintenance, especially when the new rule is more cognitively demanding than the previous one while the reward remains the same. In short, it is possible that dACC deactivation may affect either or both processes of *rule-switching* and *rule-maintenance*.

To study the interaction between the dlPFC and the dACC during cognitive control processes, we recorded the local field potentials from the dlPFC before and during the reversible deactivation of the dACC while the monkeys were performing the task. We hypothesize that changes in oscillatory activities in the dlPFC during dACC deactivation will reveal how the two regions communicate to fulfill their roles in cognitive control. Low frequency oscillations from the theta to beta range (4–30Hz) have been suggested to coordinate neural activities across brain regions to serve cognitive functions. In both the lateral PFC and the dACC, theta activities have been implicated in attention and cognitive control (Tsujimoto et al., 2006; Tsujimoto et al., 2010; Voloh et al., 2015; Voloh and Womelsdorf, 2017; Womelsdorf et al., 2010). While less studied than theta, prefrontal beta oscillations (13–30Hz) has been suggested to orchestrate cell assemblies that maintain information in short-term memory (Kopell et al., 2011), predict reaction time (Buschman et al., 2012; Ma et al., 2018), and reflect action outcome (Skoblenick et al., 2016). In between the theta and the beta bands, the alpha rhythms (9–12Hz) may help inhibit attention to irrelevant information (Buschman et al., 2012; Puig and Miller, 2015). Here, we found that cryogenic dACC deactivation impaired the animals’ task performance in a manner consistent with an impairment of *rule-maintenance*, but not *rule-switching*. This was manifested by a lower plateau in response accuracy rather than a delay in the implementation of the new rule. Correlated with this performance impairment was a reduction in the absolute difference in dlPFC fixation-period beta activities between correct and error trials, particularly for antisaccades. We also observed increased saccadic reaction time with both rules and across post-switch stages, which coincided with a reduction in task-related theta/alpha activities (5–15Hz) found for both rules in the dlPFC. Lastly, while theta-gamma phase-amplitude modulation in the dlPFC became stronger in the second halves of the sessions, during both on-and off-task periods, dACC deactivation reversed this pattern and resulted in a decrease in this cross-frequency interaction. Together our findings suggest a critical role of the dACC and its interaction with the dlPFC in the maintenance of new rules in a feedback-based rule-switching task.

## RESULTS

### dACC Deactivation Impaired Behavioral Performance on the Rule-Switch Task

To reveal the effects of cryogenic dACC (area 24c) deactivation on both behavioral performance and dlPFC (area 46/9d) activities (Figure 1A), we conducted two types of recording sessions. In the ‘cooling sessions’, after a 30-min ‘baseline epoch’ of behavioral and microelectrode array recordings, cold methanol was pumped through the cryoloops to deactivate the dACC while the recordings continued for another 30 minutes. Because the temperatures of the cryoloops took up to 4 minutes to descend to below 20°C, the initial 4 minutes of the 30-min period were excluded from further analyses, and the rest of the 30 minutes constitute the ‘cooling epoch’ (Figure 1B). Because in our design the cooling epoch took place after the baseline epoch and therefore was confounded with other variables that may affect behavior such as fatigue and reward satiation, we additionally conducted ‘sham sessions’ which alternated pseudo-randomly with cooling sessions. In sham sessions, the animals were prepared similarly and went through the first 30-min baseline epoch. For the next 30-min ‘control epoch’, the pumps were on but with no coolant running through the cryoloops. Thus, any difference in behavioral performance or neural activities between the baseline and control epochs in the sham sessions reflects the effect of epoch, while in the cooling sessions such difference reflects a combined effect of both epoch and dACC deactivation. Hence a comparison between these differences reveals any effect specific to cooling *per se*.

**Figure 1.**
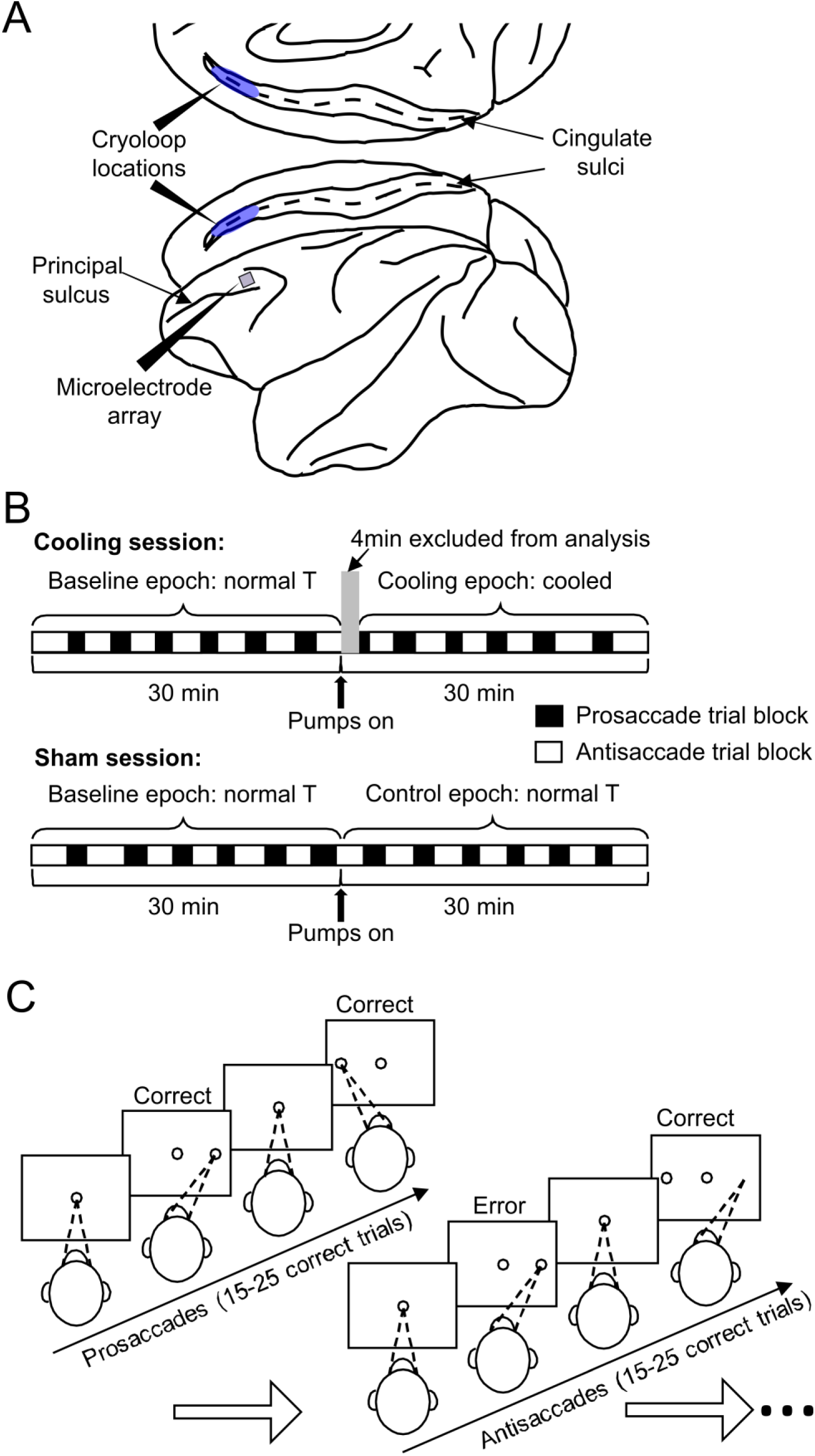
Schematics for the experimental set up, design and behavioral paradigm. **A)** The cryoloops were implanted bilaterally into the cingulate sulci (blue shades), and the microelectrode array was placed in the dorsolateral prefrontal cortex of the left hemisphere. The posterior ends of both devices were placed at the same anterior-posterior coordinate as the principal sulcus. **B)** At the beginning of a daily session, the animals were prepared similarly and performed the task for a 30-min baseline epoch before the pumps were turned on. In a cooling session, this was ensued by a 30-min cooling epoch during which chilled methanol ran through the cryoloops, which was not the case for the 30-min control epoch in sham sessions. The first 4 minutes of the cooling epochs were excluded from analyses because the temperature of the tissue was transitioning to the target range. In each session, the animal completes 12–13 trial blocks under the prosaccade (empty rectangles) and the antisaccade (filled rectangles) rules, respectively. Cooling and sham sessions were conducted in a pseudorandom order. **C)** To obtain a liquid reward, the animals were required to maintain fixation on the white central spot for a variable interval of 1.1 to 1.4s, then make a saccade toward or away from the peripheral target according to the currently applicable rule. Once a block of 15 to 25 correct trials under one rule—prosaccade in this case—the rule was switched and had to be detected by trial and error, as indicated by the first trial of the antisaccade block in the schematic.

Throughout both cooling and sham sessions, the monkeys performed the rule-switch task. In the case illustrated in Figure 1C, the animal started with the prosaccade rule. Each trial started with the onset of a white fixation dot at the center of the screen. The animal had to acquire and maintain the fixation for a random interval ranging from 1.1s to 1.4s, until the onset of a peripheral stimulus, to which a prosaccade was required for the delivery of water reward. After a random number of between 15 to 25 correct trials, the task rule was switched. In this case, when the peripheral stimulus appeared, the animal was required to generate an antisaccade away from it to the mirror location. The animals typically detected the rule switch through trial and error within approximately the first 4 trials post-switch which we labeled as the Early stage, following which they came to adopt the new rule and reached the Stable stage of performance, which we defined as the 8 trials following the Early stage (Figure 2A, top panels). For the analyses of saccadic reaction times (SRTs), we only included those from correct trials as the same SRT on different error trials may be caused by diverse erroneous processes. The SRTs also went through the Early and Stable stages as task rules switched (Figure 2A, bottom panels). Because antisaccades generally have longer SRTs than prosaccades, this switch led to an increase in SRTs in the Early stage that was sustained throughout the block of antisaccade trials (Repeated measures ANOVA, main effect of switch, *F*_1,741_ = 1889.8, p < 4.9 × 10^−324^; Figure 2A, bottom panels). At the end of the block, the rule was switched again, and the animals went through the same two stages in performance (Figure 2B, top panels). In contrast, switching from antisaccade to prosaccade involved a decrease in SRTs (*F*_1,763_ = 959.4, p < 4.9 × 10^−324^; Figure 2B, bottom panels). Each type of switches—prosaccade to antisaccade and vice versa—were repeated on average 12–13 times per session (Figure 1B). It should be noted that these switches involve more than simple reversal learning, because of the additional cognitive demand on antisaccade trials to suppress the prepotent prosaccade response. Hence, the ability to switch between rules with asymmetric cognitive demands may reflect a flexibility in deployment of cognitive resources based on the task context.

**Figure 2.**
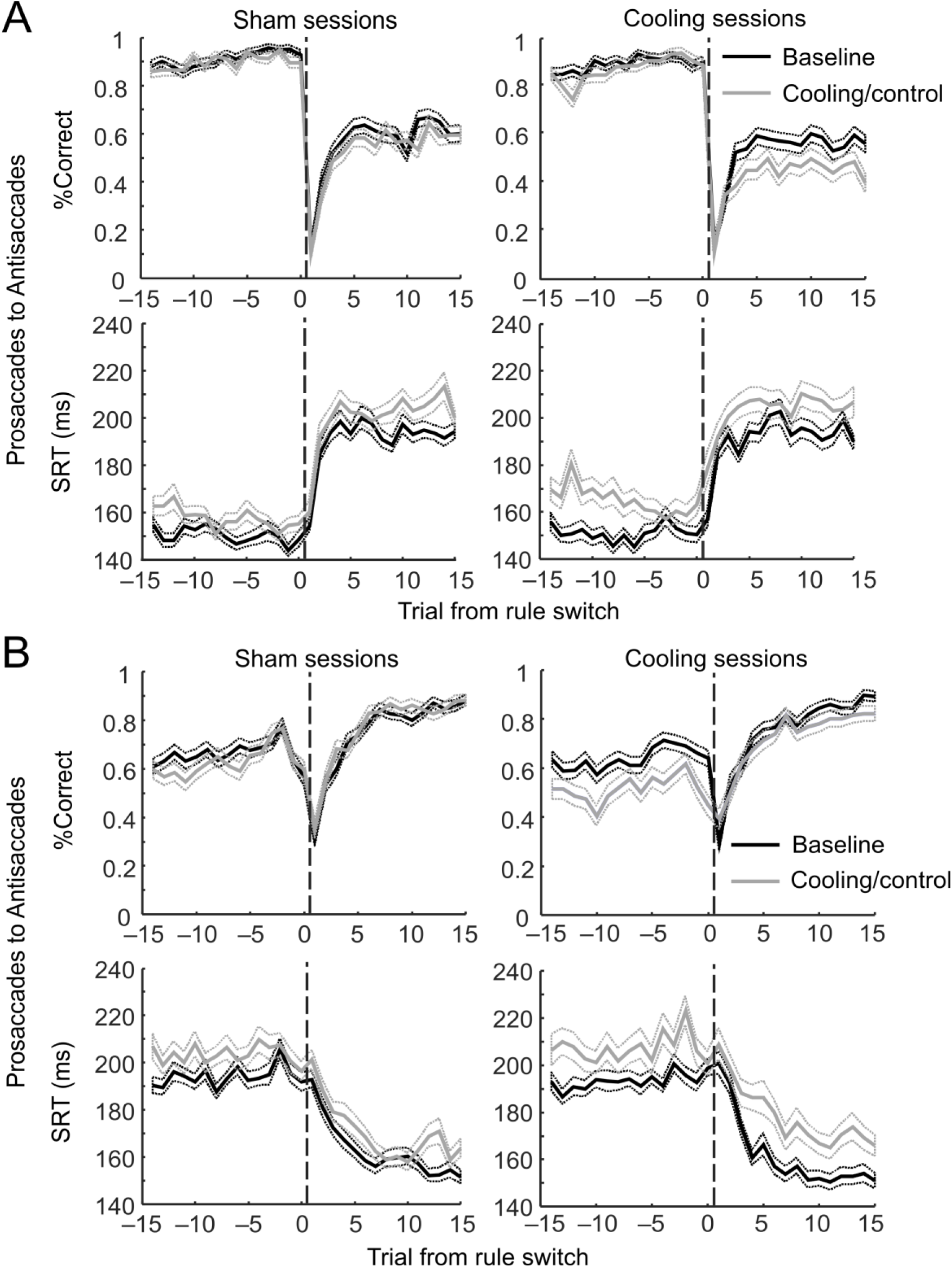
ACC deactivation impaired behavioral performance in antisaccade trials and increased the SRTs under both rules. **A)** Switches from the prosaccade to the antisaccade rule. Top panels: in both sham (left) and cooling sessions, percentages of correct trials—averaged across all trial blocks completed—dropped sharply at the first trial after the rule switches but rose quickly (the Early stage) to a plateau at approximately the 5th trial post switch. In sham sessions (top left panel), the performance remained unchanged from the baseline (black curve) to the control epoch (gray curve) on either the pre-switch prosaccade trials (to the left of the dashed line) or the post-switch antisaccade trials (to the right of the dashed line). Cooling impaired the animals’ performance in the post-switch antisaccade trials (top right panel, gray vs. black curves, to the right of the dashed line) but not in the pre-switch prosaccade trials (to the left of the dashed line). Bottom panels: cooling significantly increased the SRTs of correct responses in both the pre-switch prosaccade responses and the post-switch antisaccade responses (gray vs. black curves, bottom right panel) and more strongly than epoch alone (gray vs. black curves, bottom left panel). **B)** Switches from the antisaccade to the prosaccade rule. Top panels: similar to the top panels in **A)**, the animals’ performance went through the Early post-switch stage in which the performance rose sharply, and the Stable stage in which the performance was maintained or improved at a much slower pace. Cooling impaired the animals’ performance in the pre-switch antisaccade trials (top right panel, gray vs. black curves, to the left of the dashed line) but not in the post-switch prosaccade trials (to the right of the dashed line). No such effect was found for either rules between the baseline and the control epochs (top left panel). Bottom panels: cooling (bottom right panel) but not epoch *per se* (bottom left panel) significantly increased the SRTs in both the pre-switch antisaccade responses and the post-switch prosaccade responses. The dotted lines indicate standard error of the mean.

We examined the effect of epoch and cooling on task performance and SRT using repeated measures ANOVA on the 15 trials before and after the switches, with the ‘switch’ as the within-subject variable (pre vs. post), and epoch and session type (cooling vs. control) as the categorical factors. For the prosaccade-antisaccade switches (Figure 2A, top panels), we found a significant interaction between session type and epoch (*F*_1,745_ = 4.88, p = 0.028). Cooling impaired the animals’ performance in post-switch antisaccade trials (post hoc Tukey’s test: p = 3.2 × 10^−5^) but not in pre-switch prosaccade trials (p = 0.83); while epoch itself had no effect in sham sessions on either prosaccade (p = 0.97) or antisaccade (p = 0.084) trials. Similarly, in antisaccade-prosaccade switches, cooling (p = 7.7 × 10^−6^) but not epoch per se (p = 0.86) resulted in reduction in overall performance (epoch × cooling: F_1,764_ = 18.0, p = 2.5 × 10^−5^; Figure 2B, top panels). This effect was explained by a decrease in performance in pre-switch antisaccade trials (p = 3.2 × 10^−5^) but not post-switch prosaccade trials (p = 0.27) in the Cooling sessions. Hence, dACC deactivation impaired the animals’ response accuracy under the more challenging antisaccade rule.

For SRTs, the effect of cooling was less pronounced. While there was no main effect of session type (*F*_1,741_ = 2.63, p = 0.11) or interaction with epoch (*F*_1,741_ = 1.21, p = 0.27) on the SRTs, the pre-switch prosaccade responses were only marginally slower in sham sessions (p = 0.067) but were slowed significantly in cooling sessions (p = 0.00017). While SRTs on the post-switch antisaccade trials increased also in sham sessions (p = 0.016), this increase was intensified in cooling sessions (p = 0.0043; Figure 2A, bottom panels). In the antisaccade-prosaccade switches, we found a significant interaction between epoch and session type (*F*_1,763_ = 4.03, p = 0.045). SRTs increased from baseline in both pre-switch antisaccade trials (p = 0.0069) and post-switch prosaccade trials (p = 3.4 × 10^−5^) in cooling but not sham sessions (p = 0.063 and 0.12 respectively; Figure 2B, bottom panels). In short, dACC deactivation increased the animals’ SRTs under both rules.

In summary, dACC cooling impaired the animals’ behavioral performance on the rule-switching task. From the line plots, it appears that cooling affected the Stable stage more than the Early stage after the task switches, especially in the antisaccade trials (Figure 2A and B, top right panels), compared to the controls (top left panels).

### dACC Deactivation Specifically Affected Behavioral Performance During Rule Maintenance

Because each rule switch elicited a two-stage adaptation to the new rule (Figure 2), we subsequently carried out finer-grained analyses of the switch-related behaviors and their associated neural activities. To investigate the effects of epoch, session type and rule switch, we performed a repeated measures ANOVA for prosaccade and antisaccade trials respectively, with post-switch stage and epoch as within-subject variables and cooling as a categorical factor. Based on the distinctive stages visible in Figure 2, the rule-switch factor had two levels: Early stage—including the first 4 trials after the switch, and Stable stage—including the subsequent 8 trials, i.e. the 5^th^ to 12^th^ post-switch trials. Any effect of cooling, and not epoch alone, should take the form of an interaction between epoch and session type: data between the baseline and cooling epochs of cooling sessions would be expected to be more different than those between the two epochs of sham sessions. In prosaccade trials, neither epoch (*F*_1,58_ = 0.186, p = 0.67) nor cooling (main effect of cooling: *F*_1,58_ = 1.76, p = 0.19) affected response accuracy, although there was a marginal interaction between epoch and session type (*F*_1,58_ = 3.34, p = 0.073; Figure 3A). By contrast, in the antisaccade trials, cooling impaired the animals’ performance (main effect of cooling: *F*_1,58_ = 7.23, p = 0.0094, of epoch: *F*_1,58_ = 12.06, p = 0.00098), specifically in the Stable (p = 0.00062) stage but not the Early stage post switch (p = 0.21; Figure 3B, left bars in each panel). In sham sessions, performance on antisaccade trials did not change between the baseline and control epoch (p = 0.94 and 0.71 in Early and Stable stages respectively, Figure 3B, right bars in each panel). Thus, ACC deactivation had a deleterious effect on the animals’ ability to achieve a high level of stabilized performance specifically under the antisaccade rule.

**Figure 3.**
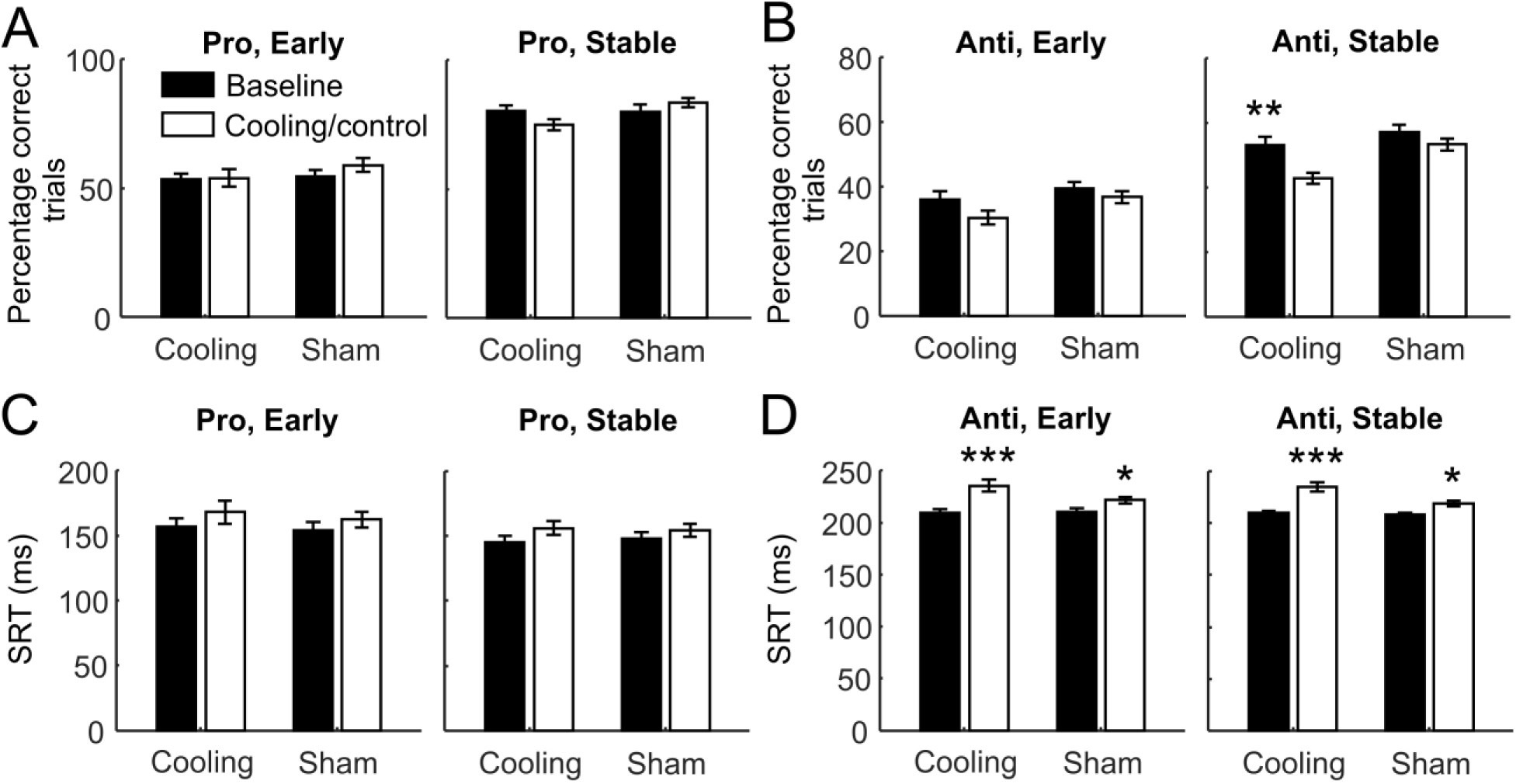
The impact of ACC deactivation on behavioral performance depended on the post-switch stage and the rule. Behavioral performance at two different post-switch stages. The Early stage included the first to the fourth trial post switch, and the Stable stage included the fifth to the twelfth trial. **A)** Performance as measured by the percentage of correct trials did not change from the baseline to the cooling or control epoch in either the Early (left panel) or the Stable stage (right panel). **B)** Cooling (left bars) but not epoch *per se* (right bars) significantly impaired the animals’ performance at the Stable stage (right panel) but not the Early stage (left panel). **C)** Cooling did not affect the SRTs of correct responses in prosaccade trials in either stage alone, but when considered together, it increased the SRTs more strongly in cooling than in control epoch compared to the baseline epochs. **D)** There was a cooling × epoch interaction: while the SRTs in both the Early (left panel) and the Stable stages (right panel) increased from the baseline to the control epochs (right bars), they increased more strongly from the baseline to the cooling epochs (left bars). The error bars indicate standard error of the mean. *p<0.05 **p<0.001 ***p<0.0005

For correct SRTs, the same analyses were conducted. Unlike the changes in performance, we found an effect of epoch (repeated-measures ANOVA, *F*_1,58_ = 23.14, p = 1.1 × 10^−5^) in prosaccade trials. Although we did not find an overall effect of cooling (*F*_1,58_ = 0.037, p = 0.85; epoch × session type: *F*_1,58_ = 1.11, p = 0.30), the SRTs from cooling sessions increased more than those in sham sessions (p = 0.00076 vs. p = 0.049) when both stages are combined. In each post-switch stage, this effect was only marginal (p = 0.055 and 0.053 in Early and Stable stages respectively; Figure 3C). In antisaccade trials, we found significant effects of cooling (main effect: *F*_1,56_ = 4.27, p = 0.043; epoch × session type: *F*_1,56_ = 8.60, p = 0.0049; Figure 3D). While the SRTs from sham sessions increased from the baseline to the control epoch (p = 0.018 and 0.023 in Early and Stable stages respectively), their increase was greater in cooling sessions (p = 0.00013 in both Early and Stable stages). To test whether this change in SRTs differed between rules and post-switch stages, we computed the difference in averaged SRT between the two epochs in each session. We found a significant interaction between rule and session type (repeated-measures ANOVA, *F*_1,56_ = 4.96, p = 0.03). While the increase in SRT was significantly greater in antisaccade than prosaccade trials in cooling sessions (p = 0.001), we found no such difference in sham sessions (p = 0.76). Thus, similarly across post-switch stages, ACC deactivation increased the animals’ SRTs under both rules, with a stronger effect on the antisaccade than the prosaccade rule.

Taken together, dACC deactivation impaired performance in the Stable stage of antisaccade trials in the rule-switching task – the effect was dependent on both the post-switch stage and the task rule. It is possible that a floor effect accounted for the lack of effect of cooling during the Early stage. For the SRTs, dACC deactivation affected antisaccade more than prosaccade trials but in a stage-independent manner for both rules. At the transition between Early and Stable stages, the performance improved significantly for each rule, epoch and session type (all comparisons for prosaccades: p = 0.00013; antisaccades: 2^nd^ epoch in cooling sessions: p = 0.00015, all other comparisons: p = 0.00013). Deactivation-specific deficits in performance became evident during the Stable stage. Although animals did not appear to have difficulty switching to the new rule, they showed deficits in maintaining it. This is potentially the result of two factors—a higher probability of committing an error after a correct response, i.e. a regression problem, or a higher frequency of consecutive errors, i.e. a perseveration problem. We therefore analyzed the correct/error pattern across sets of two consecutive trials in each post-switch stage.

We first examined the probability of committing an error after a correct response by performing a repeated-measures ANOVA with post-switch stage, epoch and rule as within-subject variables and cooling as the categorical variable. We found a significant 4-way interaction among stage, epoch, rule and cooling (*F*_1,55_ = 5.04, p = 0.029, Figure 4A,B). This effect is explained by unique increase in the percentage of errors following correct trials during cooling in the Stable stage of antisaccade trials (p = 0.033), which was not found in trials from the same stage and task rule in sham sessions (p > 0.99). Nor was this found in the prosaccade trials in the same stage (p > 0.99) or in the antisaccade trials in the Early post-switch stage (p > 0.99) during dACC deactivation. Thus, while cooling did not affect regressive errors in prosaccade trials, it increased this type of errors in antisaccade trials specifically in the Stable stage.

**Figure 4.**
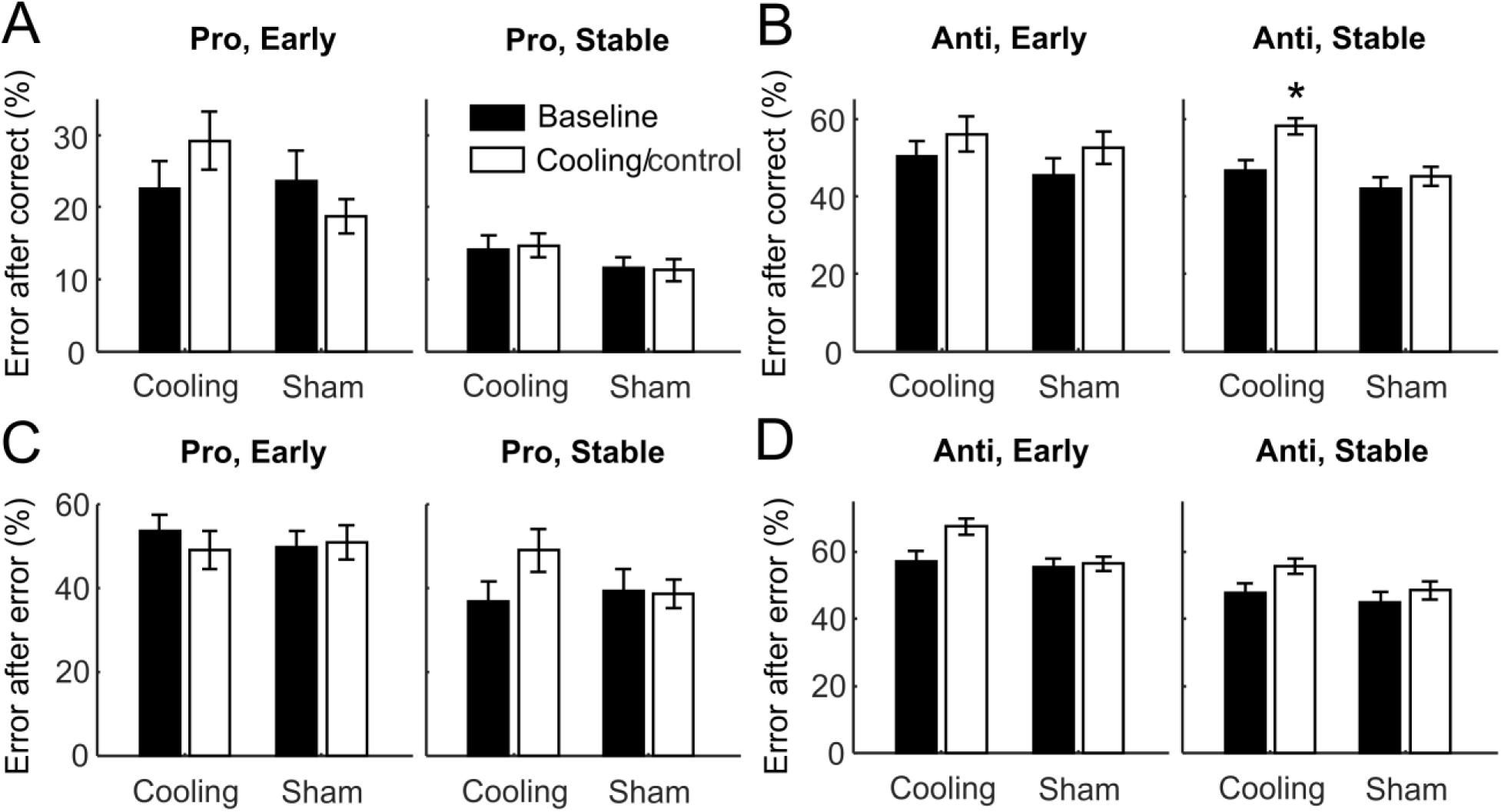
The impact of ACC deactivation on regressive and perseverative errors depended on the post-switch stage and the rule. **A)** In prosaccade trials, the percentage of errors after correct responses did not change with epoch or cooling at either post-switch stage. **B)** In antisaccade trials, such regressive errors increased with cooling during the Stable stage (right panel, left bars) but not the Early stage (left panel, left bars). Epoch *per se* did not affect this measurement (right bars in both panels). **C)** and **D)** On the percentage of errors after errors, no effect of either epoch or cooling was found for either rule or stage considered separately. When the two rules were considered together, cooling significantly increased such perseverative errors in the Stable stage (left bars in the right panels of both **C)** and **D)**) but not the Early stage (left bars in the left panels). Epoch *per se* did not have such effect (right bars in all panels).

We next investigated the probability of committing another error after an error response. There was a marginal 4-way interaction among stage, epoch, rule and cooling (*F*_1,57_ = 2.95, p = 0.091, Figure 4C,D). While not having any effect on either rule individually (prosaccades: p = 0.62, antisaccades: p = 0.98), dACC cooling significantly increased the percentage of errors following error trials in the Stable phase when trials from both rules were combined (p = 0.038), which was not seen in the Early stage (p = 0.97) or in the sham sessions (p > 0.99).

In summary, during the Stable stage, dACC deactivation lead to an increase in *regressive* errors on antisaccade trials only, and an increase in *perseverative* errors on both pro and antisaccade trials. While it is possible that the dACC supports sustained attention on the rule in our task, the rule-dependency of the cooling effect suggests a role of the dACC in the cognitive effort involved in the preparation for the more challenging antisaccades.

### dACC Deactivation Reduced Performance-Related Beta Activities in Antisaccade Trials in the Stable Stage

Given the stage- and rule-dependent effect of dACC deactivation on the rule-switching task, we investigated whether this manipulation also had performance-related effects on neural activities in the dlPFC. We focused our analysis on the fixation period, during which the animals prepared for the upcoming peripheral stimulus by retaining the current task rule. We computed the absolute difference between the power spectra on correct and error trials, then examined how this correct-error (C-E) distance changed with cooling. This was done by subtracting the C-E distance during the baseline epochs from the cooling epochs in the cooling sessions, resulting in a combined effect of both cooling and epoch (left columns, Figure 5A for Early stage and 5B for Stable stage). We also obtained the effect of epoch alone by subtracting the C-E distance during the baseline epochs from the control epochs in the sham sessions (right columns, Figure 5A and B for Early and Stable stages respectively). We then isolated the effect of cooling by performing a paired *t*-test between the pairs of power spectra for each rule at each post-switch stage (Figure 5C).

**Figure 5.**
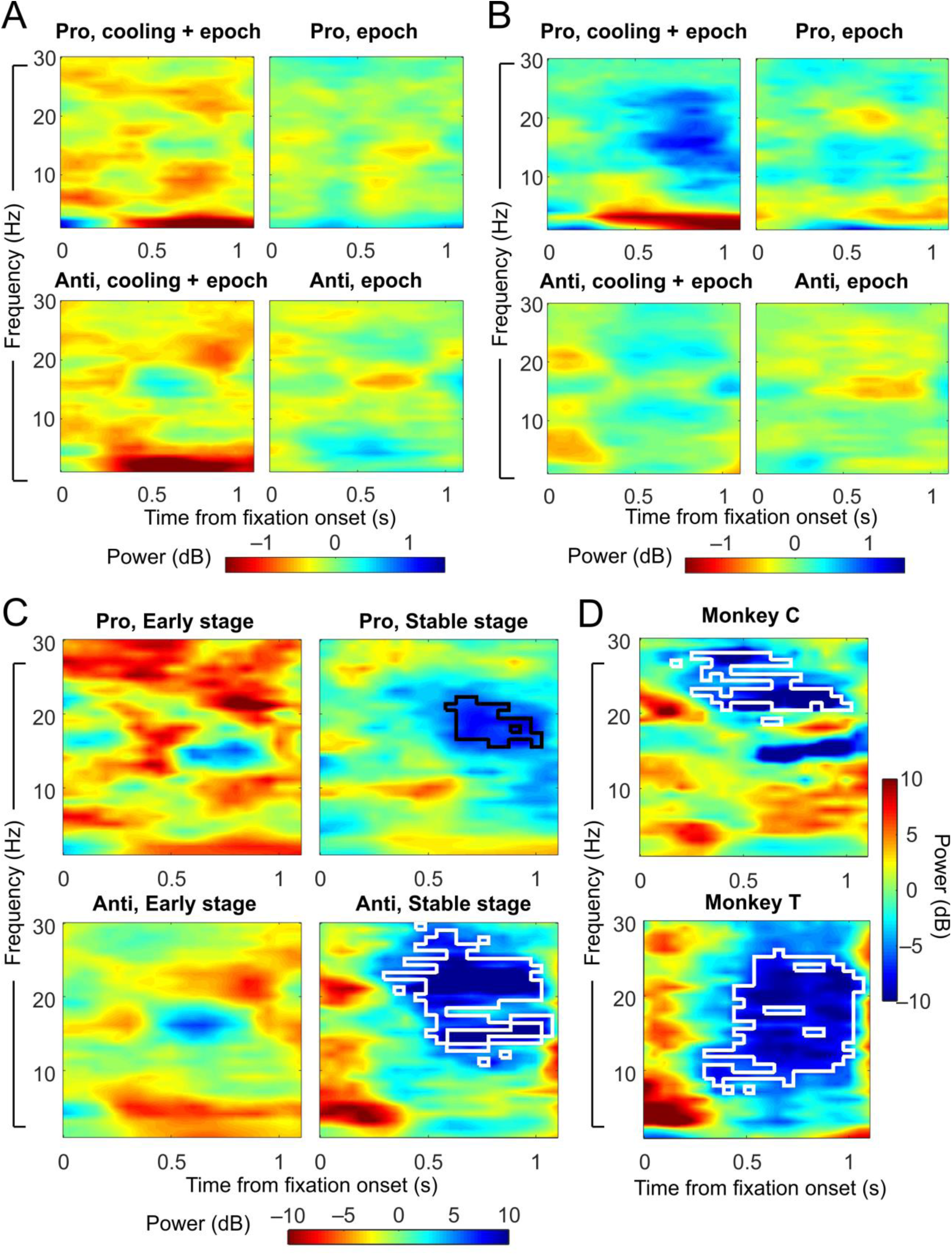
ACC deactivation reduced the correct-error difference in dlPFC theta/alpha-band LFP power in the fixation periods of antisaccade trials at the Stable stage. **A)** Change in correct-error difference in LFP powers from the baseline to the cooling epochs (left panels) or the control epochs (right panels) at the Early stage. Left panels demonstrate the effect of both cooling and epoch, while right panels show the effect of epoch only. **B)** Change in correct-error difference from the baseline to the cooling epochs (left panels) or the control epochs (right panels) at the Stable stage. **C)** *t*-statistics between the changes due to cooling and epoch combined and those due to epoch alone, which demonstrate the effect of cooling. Left panels: *t*-statistics on the effect of cooling in prosaccade (top) and antisaccade (bottom) trials at the Early post-switch stage. Right panels: *t*-statistics on the effect of cooling at the Stable stage. Cooling specifically reduced the correct-error difference in beta-band activities in antisaccade trials at the Stable stage (white contour, bottom right panel) by a cluster-based permutation test considering both rules and stages. An exploratory permutation test on the prosaccade trials at the Stable stage alone also revealed a reduction in beta-band activities (black contour, top right panel). **D)** The effect of cooling on the correct-error difference in beta-band activities in antisaccade trials at the Stable stage was found in each of the two animals. White contours mark the area of significance by a cluster-based permutation test.

Paired *t*-tests revealed visible reductions in the correct-error difference in the alpha/beta frequency band in the Stable, but not Early, post-switch stage (Figure 5C, right and left columns, respectively). This reduction reached significance in the Stable-stage for antisaccade (Figure 5C, bottom right panel, white contour) but not the prosaccade trials or in the Early stage with either rule. This effect was found in both animals (Figure 5D). To avoid the statistical pitfall of multiple comparisons, a single cluster-based permutation test was performed simultaneously for all 4 rule-by-stage groups, in which a dominant effect can overshadow other smaller effects. Hence, we also performed an exploratory permutation test for the Stable-stage prosaccade trials alone, which revealed a reduction in low beta activities associated with dACC cooling (Figure C, top right panel, black contour). This effect was also found in both monkeys. In short, dACC deactivation lead to a reduction of the difference in neural activities between the fixation periods preceding correct and error responses.

### dACC Deactivation Reduced Task-Related Theta/Alpha Oscillatory Activities

In addition to performance impairment in the antisaccade trials at the Stable stage, we also found an increase in SRTs in correct trials under both rules and across post-switch stages. We hypothesized that dACC deactivation would alter oscillatory activities in the dlPFC on these trials similarly under both rules, in a frequency band outside of the beta range, and regardless of post-switch stages. We computed and normalized the time-resolved power spectra from the fixation periods in correct trials across post-switch stages and subtracted the results of the baseline epoch from the cooling epoch in the cooling sessions to obtain a combined effect of cooling and epoch; and subtracted the baseline epoch from the control epoch in the sham sessions to obtain the effect of epoch alone (Figure 6A for prosaccades and 6B for antisaccades). We then isolated the effect of cooling by performing a paired *t*-test between the two power spectra for prosaccade (Figure 6C, left panel) and antisaccade trials (Figure 6D, left panel). In support of our hypothesis, we found a significant decrease in task-related activities in the theta/alpha range especially in the late fixation period (cluster-based non-parametric permutation test: p=0.001; blue areas with white contour, Figure 6C and D, left panels). This effect of ACC deactivation was found in each of the two monkeys for both task rules (Figure 6C and D, top and bottom right panels).

**Figure 6.**
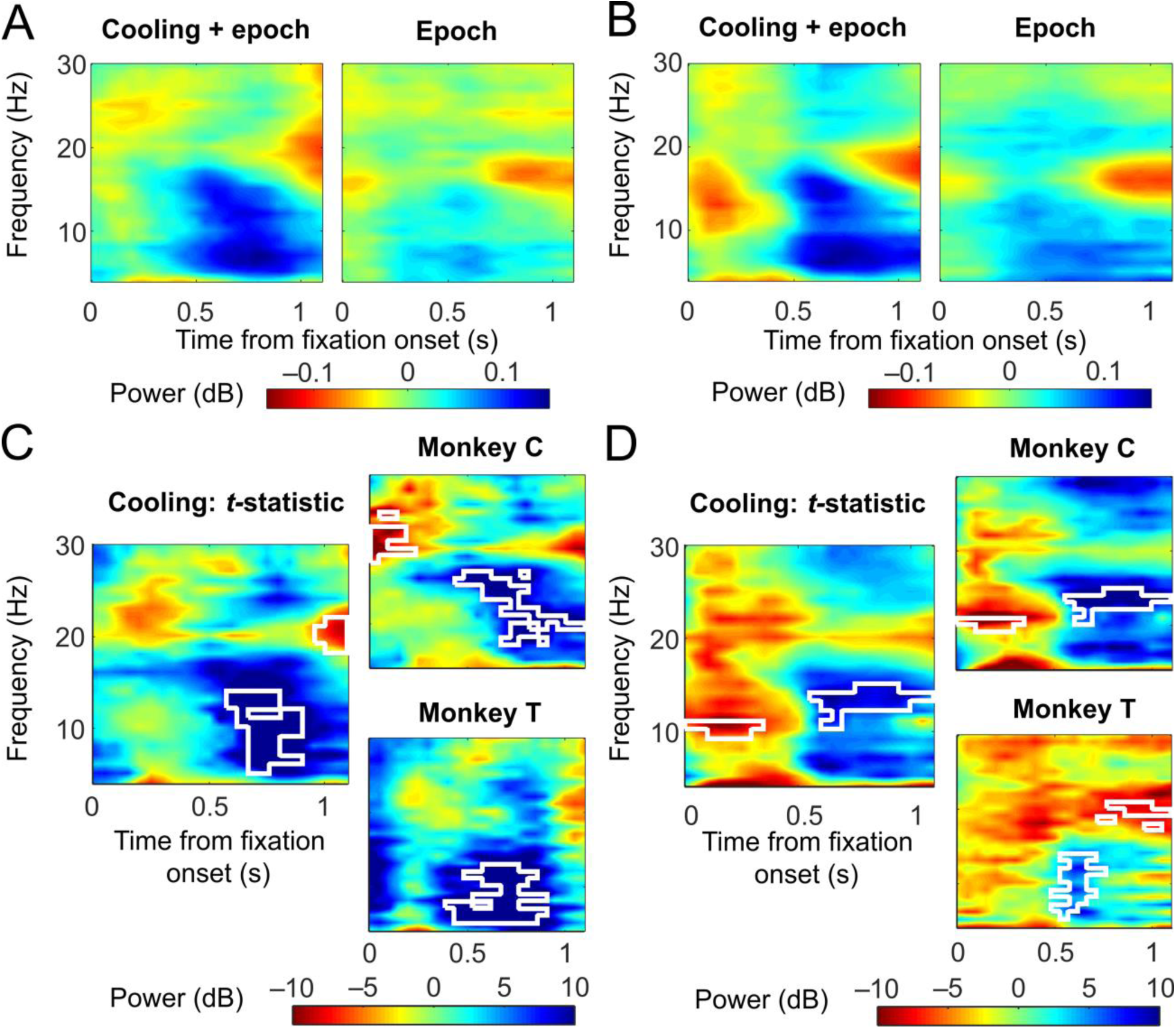
ACC deactivation reduced the fixation-period theta/alpha activities in the dlPFC during correct trials under both rules. **A)** Change in dlPFC LFP powers during prosaccade trials from the baseline to the cooling epochs (left) or to the control epochs (right). Both Early and Stable post-switch stages were included. **B)** Change in LFP powers during antisaccade trials from the baseline to the cooling epochs (left) or to the control epochs (right). **C)** *t*-statistics between the changes due to cooling and epoch combined and those due to epoch alone as shown in A), which represent the effect of cooling. Using a cluster-based permutation test, a significant reduction in theta/alpha activities was found in the second half of the fixation period (white contour, left panel), which was found in each of the two monkeys (white contours, right panels). **D)** *t*-statistics between the changes due to cooling and epoch combined and those due to epoch alone as shown in B), which reflect the effect of cooling on antisaccade trials. A significant reduction in theta/alpha activities was found in the second half of the fixation period (white contour, left panel), which was found in each of the two monkeys (white contours, right panels).

In summary, dACC deactivation reduced task-related theta/alpha oscillations in the dlPFC during correct trials, which coincided with an increase in SRTs in both rules regardless of post-switch stage. This effect was distinct from the reduction in correct-error distance we found in the beta range, which was specific to the antisaccade rule. It may be speculated that the loss of theta/alpha activities represented a decrease in motor preparation toward the end of the fixation period, which resulted in an increase in SRTs. Alternatively, given the binary nature of the task rules, it is possible that some of the correct responses were made by chance rather than under strong guidance of the rule. Hence the decrease in theta/alpha activities may reflect a weakened representation of the rule in the dlPFC, which resulted in reduced guidance and/or weakened confidence over the choice of actions, whereby prolonging SRTs.

### dACC Deactivation Impaired Theta-Gamma Phase-Amplitude Correlation

dACC-PFC theta-gamma correlation has been shown to play a role in attentional processes (Voloh et al., 2015), which are required for both successful rule switching and rule maintenance. Given that task-related theta/alpha activities were weakened by dACC deactivation, we next examined whether its interaction with gamma-band activities was also affected. We focused on how the amplitudes of gamma oscillations were distributed across different phases of low-frequency oscillations, which is independent of any change in the strength of the low-frequency activities. For completeness, we analyzed the phase-amplitude correlations between all pairings between low frequency bands including theta (4–8Hz), alpha (8–16Hz) and beta (14–28Hz), and high frequency ranges including low (30–60Hz), mid- (40–80Hz) and high gamma (80–120Hz) ranges (see Methods for more details). Only theta-low gamma phase-amplitude correlation displayed patterns of changes due to epoch and cooling consistent across the animals. During the baseline epoch in sham sessions, low gamma amplitude tended to be the greatest around the middle of a theta cycle for both prosaccade and antisaccade trials (Figure 7A, left panels, red curves). This remained unchanged in the control epochs (blue curves). Both red and blue curves had a polarity distinct from the orange curves showing the distribution of gamma amplitudes shuffled in time.

**Figure 7.**
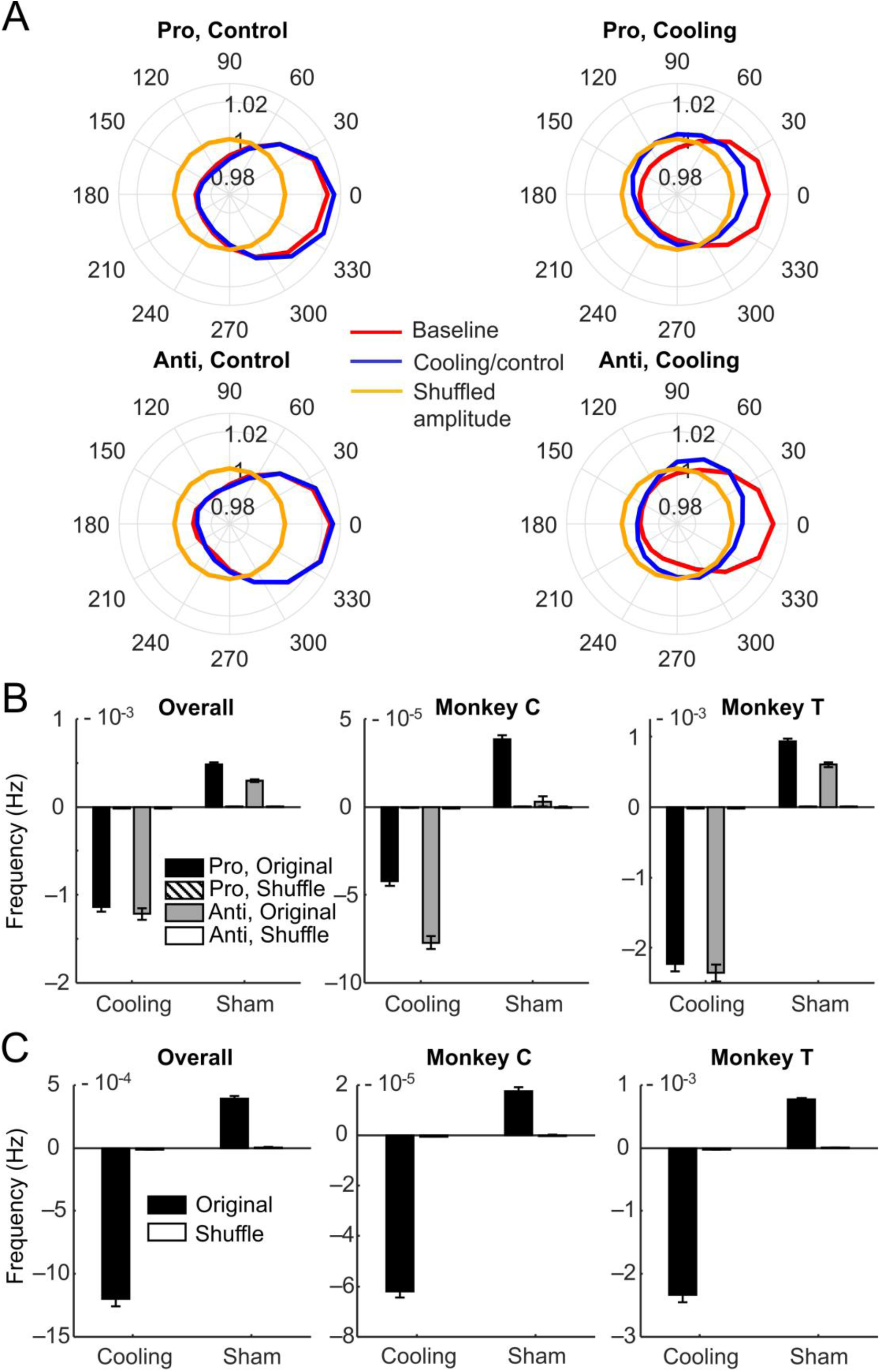
ACC deactivation reduced the theta-low gamma phase-amplitude correlations in the dlPFC. **A)** Distributions of averaged low-gamma amplitudes across different phases of a theta cycle. During the baseline epochs of both session types, low gamma amplitude tended to be the highest at the middle of a theta cycle (red curves, all panels). In the control epochs this remained the case (blue curves, left panels), but in the cooling epochs this pattern shifted (blue curves, right panels) for both rules, although some polarity remained as compared the distribution of shuffled gamma amplitudes which does not show any modulation by theta phases (orange curves, all panels). **B)** Decreases in the phase-amplitude modulation index (MI) from the baseline to the cooling epochs (left bars, all panels) and increases to the control epochs (right bars, left panel) during the fixation periods of correct trials under both the prosaccade (black bars) and antisaccade (gray bars) rules. The MI quantifies the difference between the empirical distribution of gamma amplitudes across a theta cycle from a uniform distribution. No such change was found in MIs calculated from temporally-shuffled gamma amplitudes (hatched and white bars, barely visible due to the low values with which they were associated). The opposite effects of epoch and cooling were found in each of the two monkeys (middle and right panels). **C)** The same decrease in MI from baseline to cooling epochs (left bars) and increase to control epochs (right bars) was also observed during the inter-trial intervals, in each of the two monkeys (middle and right panels) and when they were combined (left panel).

During the baseline epoch of cooling sessions, low gamma amplitude had the same tendency as in the baseline epoch of sham sessions (Figure 7A, right panels, red curves). When cooling commenced, this polarity shifted away from the middle of theta cycles and became smaller (blue curves). This comparison between session types indicates an effect of cooling on theta-low gamma phase-amplitude correlations.

To quantify the strength in theta-gamma coupling, we used the Modulation Index (MI), which is a normalized Kullback-Leibler distance between the observed distribution of mean gamma amplitudes across the theta phases and a uniform distribution with the same mean as the observed distribution. A single value of MI is generated for each pair of channels using the phases of low-frequency oscillations from one channel and the amplitudes of high-frequency activities from the other. We then calculated the amount of change in MI from the baseline to the control or cooling epoch and performed ANOVA on these changes using 3 factors: shuffling, cooling and the task rule. We found a significant interaction between session type and shuffling (*F*_1,18040_ = 1163.6, p = 3.3 × 10^−247^): while the change in MI from shuffled data did not differ between cooling and sham sessions (p > 0.99 for both rules; Figure 7B, left panel, striped and white bars) and was close to zero, with the original data, there was a significant reduction in MI in cooling sessions (original vs. shuffle: p = 6.0 × 10^−8^) and a significant increase in sham sessions (p = 6.0 × 10^−8^) for both rules (Figure 7B, left panel, black and gray bars). There was also an effect of the task rule: in sham sessions, the antisaccade rule was associated with a smaller epoch-related increase in the MI than the prosaccade rule (p = 0.0016). The greater increase in MI during the prosaccade trials may reflect a greater decrease in motivation, or it may reflect a greater increase in experience and/or confidence associated with the simpler rule. Both monkeys displayed the differential pattern of changes in MI associated with session types and with rule (Figure 7B, middle and right panels). In short, theta-gamma phase-amplitude modulation appears to depend on dACC activity, which together with the loss of task-related theta/alpha power may indicate an important role of dACC in dlPFC theta oscillations.

To determine whether this effect of dACC deactivation was task-related, we repeated this analysis on activity during the pre-trial ITIs. Since the animals were never given explicit instruction regarding the current task rule, they were required to retain this information during the ITIs. We reasoned that since any representation of the current task rule may strengthen towards the end of the fixation period to guide the forthcoming response, a greater change in MI in this epoch would indicate its task relevance; however, a lack of difference between the fixation periods and the ITIs does not completely rule out such relevance. Similar to the results from the fixation periods, there was a significant interaction between shuffling and session type (*F*_1,9020_ = 604.2, p = 3.5 × 10^−129^). While the change in MI from shuffled data did not differ between cooling and sham sessions (p = 0.98), there was significant reduction in MI in cooling sessions (p = 3.8 × 10^−9^; Figure 7C, left panel, left bars) and significant increase in sham sessions (p = 3.8 × 10^−9^; Figure 7C, left panel, right bars). This pattern held true for each of the two monkeys (Figure 7C, middle and right panels). An additional ANOVA over the effect of shuffling, cooling and fixation vs. ITI revealed no difference between the epochs (*F*_1,27064_ = 0.10, p = 0.75). In short, MI increased from baseline to control epochs, but decreased during dACC deactivation. This change was not modulated by behavior, although the increase with time was more pronounced during prosaccade trials, suggesting that theta-gamma interaction is more likely involved in the general motivational/emotional state of the animals in the alternating trial blocks rather than in the representation of rule information within individual trials.

## DISCUSSION

We set out to determine the effect of bilateral deactivation of the dACC on the performance of an uncued rule-switching task in macaque monkeys, and on dlPFC oscillatory activities associated with task performance. We observed both impairment in response accuracy and increase in saccadic reaction time, which is consistent with reports from lesion studies with human patients (Gaymard et al., 1998; Milea et al., 2003). Our findings also support a role of the dACC in presetting the saccade network especially during the cognitively demanding antisaccades, as evidenced by its stronger activation during the pre-stimulus preparatory period before antisaccades than prosaccades in human fMRI studies (Brown et al., 2007; Ford et al., 2005).

Specifically, we found that dACC deactivation impaired the animals’ performance on the more cognitively demanding antisaccade trials, especially during the Stable post-switch stage rather than the initial transition to the antisaccade rule (Figures 2,3). Correlated with this finding was a reduction in the difference in beta band activities between correct and error trials on antisaccade trials during the Stable stage only (Figure 5). This correlation between changes in beta activities and performance has been reported in several studies (Babapoor-Farrokhran et al., 2017; Buschman et al., 2012; Skoblenick et al., 2016). In the current task, the rule information must be maintained across trials rather than being cleared from working memory at the end of each trial. In a task where the prosaccade and antisaccade rules were cued in each trial and the trial types were randomly interleaved, beta activities were weaker during the task epochs than during the ITIs (Ma et al., 2018). This is consistent with a role of prefrontal beta rhythm in gating information coding and inhibiting potential interference (Lundqvist et al., 2018; Lundqvist et al., 2016). We reasoned that our task requires an optimal level of prefrontal beta activity for the maintenance or (re)activation of the current rule (Kopell et al., 2011; Spitzer and Haegens, 2017; Spitzer et al., 2010; Tallon-Baudry et al., 1999) as well as the suppression of the alternative rule, which marks the difference between correct and error trials. A reduction in this difference during dACC deactivation may lead to the animals’ inability to maintain a high level of performance (Figures 2,3). This reasoning is supported by the marked increase in regressive errors specifically in the antisaccade trials in the Stable stage, during which they reverted to the prosaccade rule right after a correct antisaccade response and after having shifted towards the new rule (Figure 4). The deficit may not be specific to the motor preparation for antisaccades, as the same increase in regressive errors was observed in a dACC lesion study on reversal learning using hand movements (Kennerley et al., 2006). In summary, the best behavioral performance may require an optimal level of beta activities in the LPFC, which is not a localized phenomenon but relies on an intact network that includes the dACC.

While performance remained the same in prosaccade trials, dACC deactivation increased reaction times under both rules and to the same extent in both Early and Stable post-switch stages (Figure 3). Correlated with this behavioral change was the reduction in task-related theta/alpha band activities (5–15Hz) in the dlPFC during the second half of the fixation periods, which was found for both rules (Figure 6). The dACC is the primary source of ‘frontal midline theta’ activities in EEG studies, which are believed to reflect signals of cognitive control and negative affect (Cavanagh and Shackman, 2015). Additionally, dACC theta was found to carry rule information prior to the onset of the stimulus that determines the direction of the saccade (Womelsdorf et al., 2010). Because task information can be transferred from the dACC to the lateral PFC through theta activities (Smith et al., 2015), the reduction in task-related theta/alpha activities in the dlPFC may reflect a loss of dACC input on either cognitive control or rule representation. Given that the decrease in theta/alpha activities coincided with longer reaction times but not necessarily errors, and that the animals improved significantly from the Early to the Stable stage even with dACC deactivation, the rule information must be preserved in other regions in the network such as the dlPFC, although its guidance over the action may be weakened. Alternatively, the loss in dlPFC theta/alpha activities may be associated with a decrease in motor preparation, which led to increased SRTs. Given the asymmetrical increase in the SRTs of antisaccades compared to prosaccades, such a preparatory function must also have a cognitive control component, which is consistent with the role proposed for both the dlPFC and the dACC in the extended saccade network (Brown et al., 2007).

While a task-related change in alpha activities is not found as often as in theta rhythms, our finding of dACC deactivation-induced reduction in amplitude ranges from 5 to 15Hz, encompassing both frequency bands. This range is consistent with findings from an intracranial recording study in humans which found a positive correlation between mental arithmetic performance and LPFC activities between 4 and 12Hz (Halgren et al., 2002). It is possible that alpha plays a different role from theta activities in our task as well as mental arithmetic, e.g. inhibiting the irrelevant rule as opposed to sustaining attention on the relevant rule, as previously suggested (Buschman et al., 2012; Schroeder et al., 2018).

Tasks demanding cognitive control engages a network involving multiple brain regions, each of which partakes in this process through oscillatory and spiking activities occurring at coordinated timing, e.g. through phase synchronization or cross-frequency coupling (Varela et al., 2001; Womelsdorf and Fries, 2007). Indeed, several human EEG studies have demonstrated a correlation between the modulation of gamma amplitude by theta phase in the frontal cortices and working memory processes (Canolty and Knight, 2010; Lisman and Jensen, 2013; Rajji et al., 2017). Weakening in this cross-frequency interaction has been correlated with the severity of cognitive impairment and dementia in patients (Goodman et al., 2018). Here we found that the phase-amplitude modulation between theta (4–8Hz) and low gamma (30–60Hz) from different recording sites in the dlPFC increased during the session (Figure 7). Adding dACC deactivation to the effect of epoch completely reversed the pattern: the phase-amplitude modulation now decreased (Figure 7). This could not be explained by any overall change in gamma amplitude, which was scaled to range between 0 and 1 for each frequency and each channel. Given that the analysis used correct trials only, it is possible that the increase in theta-gamma interaction with epoch reflected an increase in cognitive effort to compensate for fatigue and reduced motivation.

The same pattern was present during both fixation periods and the ITIs. Given that the animals must maintain the current rule throughout the session and to update it whenever necessary, we cannot rule out that the possibility that the effect found during the ITIs was still task-related, even though no decision regarding the saccadic response was made then. Since dACC theta can propagate feedback information to the LPFC (Smith et al., 2015) and modulates gamma activities in the LPFC during attentional processes (Voloh et al., 2015), it is not unexpected that dACC deactivation disrupted the theta-gamma interaction within the LPFC (Figure 7). Although the animals remained capable of updating the task rules based on feedback (Figure 2,3), they did commit more perseverative errors in the Stable post-switch stage during dACC deactivation (Figure 4), which may reflect a subtle deficit in feedback-related cognitive flexibility.

In summary, using an uncued rule-switching task, we found that dACC deactivation impaired performance maintenance after shifting to the new rule rather than slowing the rule switch per se. While our findings do not preclude the role of the dACC in error monitoring and feedback-related processing, they strongly support its function in maintaining optimal performance especially in a task with greater cognitive demands (Chong et al., 2017; Croxson et al., 2009; Hosking et al., 2014; Klein-Flugge et al., 2016; Parvizi et al., 2013; Rudebeck et al., 2006; Vassena et al., 2014; Walton et al., 2003). Importantly, our finding indicates that this role of the dACC may be accomplished partly through its influence on the task-related beta activities in the dlPFC. dACC lesion studies on reversal learning have also found a similar post-switch deficit, although the maintenance of performance on the two hand movements—which were equally cognitively demanding—were similarly impaired (Chudasama et al., 2013; Rudebeck et al., 2006). Authors from these studies suggested that the dACC was required to integrate history of both reward and errors to guide effective choice—a hypothesis that is supported by our finding that reaction times increased in both trial types with dACC deactivation, even though the performance on prosaccade trials remained relatively intact. Our results also indicate that the cognitive control signal which guides effective responses with rule representations and/or the history of outcomes may be achieved through a theta/alpha input from the dACC to the dlPFC (Smith et al., 2015). Taken together, our findings suggest that feedback-based cognitive flexibility and other aspects of cognitive control reflect the concerted effort of an extended network, in which the dACC may broadcast different signals through different oscillatory rhythms.

## MATERIALS AND METHODS

Two adult male macaque monkeys C *(Macaca mulatta)* and T *(Macaca fascicularis)*, weighing 6.5 and 9.5 kg respectively, were used in the study. After the initial chair training, they were implanted with a plastic head restraint for head-fixation during training. Once recovered, they were trained on the uncued rule-switch task. A second surgery was then conducted to implant the cryoloops and the microelectrode array once they fully acquired the task. After recovery, they were retrained before testing started.

### Surgical Procedures

Each surgery was performed aseptically, with the animals’ physiological parameters continuously monitored and frequently recorded by an experienced veterinary technician. Following each surgery, the animals received analgesics and antibiotics and were monitored by a university veterinarian. In the first surgery, the monkeys were implanted with a plastic head restraint, secured to the skull using bone screws and dental acrylic using previously described aseptic surgical procedures (DeSouza and Everling, 2004). Upon recovery from the surgery, they were trained to perform the uncued prosaccade and antisaccade switch tasks. Once trained, they underwent a second surgery in which stainless steel cryoloops (8−10 × 3 mm) were implanted bilaterally into the anterior cingulate sulci. The posterior ends of the cryoloops were placed at the same anterior-posterior coordinates as the posterior ends of the principal sulci, such that the cryoloops targeted the dACC (area 24c). Neurons with task-selective activity for prosaccades and antisaccades have been found in this area of the dACC (Johnston et al., 2007). The technical details of the cryoloop surgery and deactivation method have been previously described (Lomber and Payne, 2000; Lomber et al., 1999).

In the same surgery, each of the animals was implanted with a 48-channel Utah microelectrode array (Blackrock Microsystems LLC, Salt Lake City, UT, USA). The initial craniotomy for cryoloop placement was extended using rongeurs until the arcuate and principal sulci could be visually identified. The meninges were carefully removed, the array was placed at the center of area 9/46d in the left hemisphere and inserted with a pneumatically-actuated impulse inserter (Blackrock Microsystems LLC, Salt Lake City, UT, USA). A layer of Gel foam was then placed over the exposed brain tissue and dura mater and covered with silicone sealant before dental acrylic was applied to seal the craniotomy. The reference wires were secured underneath the skull and above dura mater. The array was connected by a bundle of fine wires to an Omnetics connector (Omnetics Connector Corporation, Fridley, MN, USA), which was then secured to the surface of the skull with dental cement at a location posterior to the implantation site and was protected by a PEEK chamber covered with a cap. The wire bundle was also secured and covered with dental acrylic.

All procedures were conducted in accordance with the Canadian Council on Animal Care Policy on the Use of Laboratory Animals, and a protocol approved by the Animal Care Committee of the University of Western Ontario Council on Animal Care.

### Behavioral Task

Monkeys were trained to perform and switch between uncued prosaccades and antisaccades (Figure 1C). Each trial began with the presentation of central white fixation spot. Monkeys were required to fixate within a 0.5° × 0.5° window surrounding this spot for a duration of 1.1s to 1.4s at the beginning of each trial. Subsequently, the fixation spot was extinguished and a peripheral white stimulus (0.15°) was presented pseudorandomly with equal probability at an eccentricity of 8° to the left or to the right. To receive a liquid reward, monkeys were required to generate a saccade within 500ms toward the stimulus on prosaccade trials and away from the stimulus on antisaccade trials, within a 5° × 5° window. After 15–25 correct trials, the task rule switched from prosaccades to antisaccades or vice versa without any explicit signal to the subjects. Consequently, rule switches were guided by trial and error based on the presence or absence of reward after each trial. Several task rule switches were completed within each session.

### Reversible Cryogenic Deactivation and Data Acquisition

To deactivate the dACC, chilled methanol was pumped through a cryoloop with Teflon tubing, which passed through a methanol and dry ice (~ −80°C) bath. Methanol that had passed through the cryoloop was returned to the same reservoir from which it came. Evoked neural activity in underlying cortical tissue is absent when cortical tissue is cooled to below 20°C (Adey, 1974; Jasper et al., 1970; Lomber and Payne, 2000).

Cryoloop temperature was monitored by an attached microthermocouple and maintained at 1–3°C to deactivate as large an area of cortical tissue as possible, while avoiding potentially damaging subzero temperatures at the cortical surface (Lomber et al., 1999). For monkey T, we had to maintain the cryoloop temperature substantially higher at 14–15°C because the monkey stopped performing the task at lower temperatures. Given that the effective spread of cooling is restricted to about 2 mm on either side of the cryoloop (Payne and Lomber, 1999) and that each cryoloop measured 8–10 × 3 × 2 mm, the volume of deactivated cortex can be approximated by the volume of a box with dimensions 12–14 × 7 × 4 mm, or 336–392 mm^3^. Thus, cooling of the cryoloops affected the dorsal and ventral banks of the anterior cingulate sulci, corresponding to area 24c (Figure 1A). Details of the cryoloop procedure have been described previously (Lomber et al., 1999).

In each cooling session, the animals performed the task for 30 minutes without cooling— the baseline epoch, followed by a 30-min cooling epoch, allowing for the collection of sufficient behavioral and neural data under both conditions. At the end of the 30^th^ minute, the pumps were turned on to start cooling. The first 4 min after the onset of the pumps were excluded from all data analysis to ensure that the cortical tissue adjacent to the cryoloops was cooled below 20°C and that neurons were deactivated (Figure 1B). In addition, both monkeys performed sham sessions that also consisted of two consecutive 30-minute epochs to control for the effects of time and fatigue over the course of a session. During the second 30-min ‘control epoch’, the pumps were turned on, but no methanol ran through the cryoloops and cortical tissue remained active (Figure 1B). Monkey C completed 17 cooling and 17 sham sessions, while monkey T performed 13 cooling and 14 sham sessions.

Throughout each cooling or sham session, eye movements were recorded at 1000 Hz with high-speed infrared video eye tracking (Eyelink 1000, Kanata, ON, Canada), and the timing of behavioral events were controlled and recorded by the Cortex real-time behavioral control and data acquisition system (NIMH, Bethesda, MD, USA). Both eye tracking and behavioral event timestamps were also sent to and recorded by a Plexon Multichannel Acquisition Processor (MAP) system (Plexon Inc., Dallas, TX, USA), which acquired local field potentials and spike trains at 1kHz from the Utah array. The MAP system synchronizes and combines different types of data into a single file.

### Data Analysis

#### Behavioral performance

All analyses were performed using custom Matlab (The Mathworks Inc., Natick, MA) code. Saccade onset was identified as the time at which saccade velocity exceeded 30°/s and saccade offset was identified as the time at which saccade velocity fell below 30°/s. SRT was defined as the time from stimulus onset to saccade onset. Trials with no fixation, broken fixation prior to peripheral stimulus onset, or with saccadic reaction times (SRTs) below 80ms or above 500ms were excluded from further analyses. Included in the analyses were the direction errors, in which a prosaccade is erroneously made in place of an antisaccade or vice versa.

Initial analysis included all trials (Figure 3), all correct trials (Figure 4) or 15 pre-switch and 15 post-switch trials from each trial block containing a pro-to-anti or anti-to-pro rule switch (Fig. 2). In later analyses, the post-switch trials were broken down to ‘Early stage’ which includes the 4 trials following the change in task rule, and ‘Stable stage’ which includes the 5^th^ to 12^th^ trials following the rule switch. Comparing the two stages provided insight into how the animals behaviour progressed from the initial rule change, signaled by omission of reinforcement, to the establishment of the new rule.

#### Preprocessing and power spectra

LFP data were analyzed in MATLAB (MathWorks, Naticks, MA, USA) using the FieldTrip toolbox (http://fieldtrip.fcdonders.nl/) developed at the Donders Institute for Brain, Cognition and Behaviour (Oostenveld et al., 2011). The recorded LFPs were low-pass filtered at 150Hz and line noise was removed at 60Hz and 120Hz using a discrete Fourier transform. Electrical noise introduced into the neural recordings by the operation of the pumps used to run methanol through the cryoloops was removed using the *chunkwiseDeline* function (https://xcorr.net/2011/10/03/removing-line-noise-from-lfps-wideband-signals/) by Patrick Minealt. This function is suitable for mechanical noise with a fixed shape in the time domain. It detects and describes the shape of the noise in the time domain with a family of exponential functions, and then subtracts it from the signal. The chunkwise delining method is preferable to the use of a notch filter since it preserves the physiological signals that may occur at the same frequency as the noise.

The continuous signals were then divided into discrete trials based on event timestamps. The first 1.1s from all trials including a saccade were included in subsequent analyses, and the 1.1s preceding the acquisition of fixation in each trial was used as inter-trial interval for normalization of LFP powers. For the time-frequency presentation of LFP power, the data was processed using a multitaper method with a discrete prolate spheroidal sequence (DPSS) taper set, using a 0.667s window in time and a 4.5Hz window in frequency for power spectra from 1 to 60Hz, and 0.333s and 12Hz windows for power spectra from 60 to 150Hz. To compare LFP between channels and animals and to reduce variability, we used decibel normalization for each channel at each frequency (Herrmann et al., 2014):

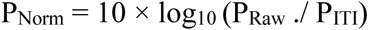
where P_Raw_ refers to the LFP powers during the fixation period and P_ITI_ refers to powers during inter-trial intervals prior to fixation.

#### Phase-amplitude correlation

We visualized the relationship between the phase of low-frequency LFPs and the amplitude of high-frequency LFPs using polar plots, and quantified the correlation using Tort’s modulation index (MI) (Tort et al., 2010; Voloh et al., 2015). Raw LFP signals, after noise removal (see previous section), were bandpass filtered using a fourth-order, two-pass Butterworth filter. The filters were defined as ± 1/3 of the center frequency, which was found to facilitate the detection of cross-frequency interactions. We defined the low-frequency bands as 4–8Hz (i.e. 6 ± 2Hz), 8–16Hz (i.e. 12 ± 4Hz) and 14–28Hz (i.e. 21 ± 7Hz). For high-frequency ranges we chose 30–60Hz (i.e. 45 ± 15Hz), 40–80Hz (i.e. 60 ± 20Hz) and 60–120Hz (i.e. 90 ± 30Hz). We analyzed the phase-amplitude correlations from all 9 low-high frequency band pairs, for each task rule, each epoch (i.e. baseline or cooling/control epoch) and each session type (i.e. cooling or control). Within each of these categories, the first 1.1s of the fixation period from all trials were concatenated into one signal, which was Hilbert transformed. From the results we obtained the phase and amplitude of each channel at each time bin in each frequency band. We analyzed only the correlations between phases and amplitudes recorded from different channels to avoid spurious coupling due to nonstationary inputs (Aru et al., 2015). For each low frequency band, the phase data were divided into *N =* 16 bins, and the amplitude data from each high frequency band corresponding to each bin *i* was averaged to yield *P*(*i*). We then normalized these mean amplitudes for each channel pair against their mean, before averaging across all channels, to give rise to polar distributions plotted in Figure 7A. We also calculated a control distribution for each rule and session type for the cooling/control epochs only. This was done by shuffling the amplitudes within each channel, which destroyed any correlation between the phases and the amplitudes without changing their values. The shuffled control distributions are also plotted in Figure 7A.

The Tort’s Modulation Index (MI) is based on the Kullback-Leibler (KL) divergence—also known as Shannon entropy or relative entropy—between two distributions: the actual distribution *P* of amplitudes across the bins and a uniform distribution *Q* of amplitudes with the same average as the observed distribution. We performed this analysis according to methods described in Voloh *et al*. (2015). Specifically, the KL divergence (D) is calculated as follows:

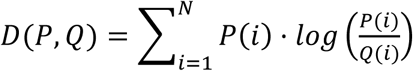

The MI is then calculated as the normalized KL divergence between the two distributions:

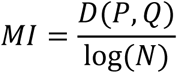

The greater the MI, the farther from uniform is the actual distribution of high-frequency oscillatory amplitudes across the phase bins of low-frequency oscillations. In other words, the MI quantifies the level of phase-amplitude cross-frequency correlation.

#### Experimental design and statistical analyses

The animals’ behavioral performance, calculated as percentage correct in a 15-trial block, was analyzed with repeated measures ANOVA, with session type (cooling or control) as the categorical factor, epoch (baseline or cooling/control) and switch condition (pre- or post-switch) as repeated measures (Figure 2). The pre-switch performance was calculated from the last 15 trials before a rule-switch, and the post-switch performance was calculated from first 15 trials after the switch. The sample size was the number of trial blocks, pooled across sessions, ranging from 152 to 225 depending on the session type, epoch or switch condition. The SRTs were analyzed similarly, with one averaged SRT used for each trial block (Figure 2). To quantify behaviors from different post-switch stages, since there were only 4 or 8 trials in each stage in a given trial block, we calculated the response accuracy or averaged SRTs based on all trials from that stage from a session, resulting in a sample size equal to session number (N = 30 and 31 for cooling and control, respectively; Figures 3 and 4).

Because the microelectrodes remained in the same locations in the brain throughout the experiment, for each animal, for the neural analyses we pooled trials recorded from different sessions and computed the power spectra using all trials with a completed response in each rule, epoch and session type. All neural analyses focused on the fixation period in each trial, which was when the animals had to prepare for the upcoming peripheral stimulus by retaining the current task rule. For statistical comparison between power spectra, we used a nonparametric cluster-based method (Maris and Oostenveld, 2007), the sample size being the number of channels (n = 96, with 48 from each animal). First, a map of t-statistics was calculated between the two power spectra (i.e. power values at each time bin and frequency). Then the significance level was determined from a distribution of t-statistics generated by 5000 iterations of pooling and random splitting of the data. Original t-statistics that were greater than 99.9% of the generated distribution were considered significant. Significant t-statistics were then clustered using the *clusterdata* function in MATLAB.

## ACKNOWLEDGEMENT

This research was supported by a Foundation grant from the Canadian Institutes of Health Research (FRN 148365) to S.E. and a postdoctoral fellowship from the Canadian Institutes of Health Research to L.M.

## AUTHOR CONTRIBUTIONS

Conceptualization, K.J., S.G.L. and S.E.; Methodology, L.M., J.L.C., K.J., S.G.L. and S.E.; Investigation, L.M., J. L.C., K.J. and S.E.; Writing – Original Draft, L.M., J.L.C.; Writing – Review & Editing, L.M., J. L.C., K.J., S.G.L. and S.E.; Funding Acquisition, S.E.; Resources, S.E.; Supervision, S.E.

